# Simultaneous Modeling of Reaction Times and Brain Dynamics in a Spatial Cuing Task

**DOI:** 10.1101/2020.11.16.384198

**Authors:** Simon R. Steinkamp, Gereon R. Fink, Simone Vossel, Ralph Weidner

## Abstract

Understanding how brain activity translates into behavior is a grand challenge in neuroscientific research. Simultaneous computational modeling of both measures offers to address this question. The extension of the dynamic causal modeling (DCM) framework for BOLD responses to behavior (bDCM) constitutes such a modeling approach. However, only very few studies have employed and evaluated bDCM, and its application has been restricted to binary behavioral responses, limiting more general statements about its validity.

This study used bDCM to model reaction times in a spatial attention task, which involved two separate runs with either horizontal or vertical stimulus configurations. We recorded fMRI data and reaction times (n=29) and compared bDCM to classical DCM and a behavioral Rescorla-Wagner model using goodness of fit-statistics and machine learning methods.

Data showed that bDCM performed equally well as classical DCM when modeling BOLD responses and better than the Rescorla Wagner model when modeling reaction times. Notably, only using bDCM’s parameters enabled classification of the horizontal and vertical runs suggesting that bDCM seems to be more sensitive than the other models. Although our data also revealed practical limitations of the current bDCM approach that warrant further investigation, we conclude that bDCM constitutes a promising method for investigating the link between brain activity and behavior.

## Introduction

Computational modeling can deepen our understanding of how the brain processes information and produces overt behavior. In the field of psychology, computational modeling has a long history in describing and explaining behavioral concepts. For example, reinforcement learning algorithms have been used to explain classical conditioning (Rescorla et al., 1972), drift-diffusion models have been used to model reaction times in decision-making tasks (Ratcliff, 1978), and race models have been used as theoretical formulations of visual-spatial attention (Bundesen, 1990). Similarly, different computational modeling approaches have been employed in the fields of neuroscience and neuroimaging. For example, generative graphical models of brain connectivity describing blood oxygenation level-dependent (BOLD) amplitudes in response to experimental inputs can be estimated using dynamic causal modeling (Friston et al., 2003, 2017), and multivariate temporal response functions have been used to model ongoing sensory stimulation, like speech, in electrophysiological recordings (Crosse et al., 2016).

Although computational models are very prominent in the two fields, behavioral and neural responses are mostly treated separately (Turner et al., 2017). However, a combined modeling approach could provide us with deeper insights into the neural processes and the emergence of behavior. Here, different approaches have been proposed: One possibility is to correlate the parameters of neural and behavioral models to describe how the different measures are related across different participants (Vossel et al., 2016). Alternatively, in model-based fMRI, the behavioral computational model’s outputs (or hidden states) are used as a factor in a classical GLM analysis. One such factor could be a participant’s perceived cue validity in a probabilistic spatial cueing task, which can be recovered from reaction times (e.g., Dombert et al., 2016). Leveraging the theory-driven performance of cognitive models allowed to determine more specific brain activation patterns of cognitive processes than by using non-specific measures such as reaction times (Turner et al., 2017). A third option is a joint modeling approach (Turner et al. 2017). Here, an overarching set of parameters is used to describe both brain activity and behavior. An example is a study by Nunez et al. (2015), where the drift-diffusion model parameters were constrained with task-based brain activity, incorporating the covariation between reaction times and neural-activity on a trial-by-trial basis.

Although these approaches are tremendously useful, none of them employs an integrative model describing the generation of brain activity and behavior, allowing us to investigate the hidden processes behind the two measurements directly. Rigoux and Daunizeau (2015) have provided such a framework, where DCM is extended by an additional output function to describe behavioral responses (behavioral DCM, bDCM). This simultaneous modeling does have not only high descriptive power but also allows thorough diagnostics of the model. For example, by disabling specific nodes in the network (i.e., artificial lesions), conclusions can be drawn about the contribution or necessity of different brain regions to the emergence of behavioral patterns. So far – to our knowledge – bDCM has been applied to a larger datasets in one study, which modeled binary choices in an economic decision making task (Shaw et al., 2019).

In the current study, we show that bDCM can be extended to continuous measures (i.e., reaction times). Furthermore, we provide a direct comparison between bDCM and classical DCM and between bDCM and an adjusted version of the Rescorla-Wagner model (Rescorla et al., 1972; Vossel, Mathys, et al., 2014).

As a testing ground, we modeled the effects of attentional reorientation along the horizontal and vertical meridians in a spatial cueing-paradigm, where participants had to report the orientation of a pre-cued Gabor patch. In trials in which invalid cues indicated an incorrect location of the target Gabor patch (20 % of the trials), participants had to reorient their attention to the opposite location (Posner, 1980). This paradigm has been found to elicit reliable reaction time differences between invalid and valid trials, both on the individual and the group level (Hedge et al., 2017). Additionally, it has been shown that the internal representation of cue-validity can be modeled using the Rescorla-Wagner model as a generative model of reaction times (Mengotti et al., 2017; Rescorla et al., 1972; Vossel, Mathys, et al., 2014).

Besides the reliable behavioral effects, the cortical networks involved in this task have been characterized by multiple studies. We have previously analyzed the present dataset using classical DCM (Steinkamp et al., 2020), which has also been used in similar cueing paradigms (c.f., Vossel et al., 2012). Moreover, studies in patients with stroke-induced lesions have revealed brain regions that are critically involved in spatial cueing-tasks (Corbetta & Shulman, 2011; Malherbe et al., 2018; Posner et al., 1984). It is well established that the orientation of visual-spatial attention is mediated by a dorsal fronto-parietal attention network consisting of the intraparietal sulci (IPS) and the frontal eye fields (FEF). This network interacts with a ventral fronto-parietal attention network of ventral frontal cortex, and the temporoparietal junction (TPJ) when a sudden reorientation of attention is necessary (Corbetta et al., 2005; Corbetta & Shulman, 2011).

In addition to comparing bDCM, which simultaneously models behavior and brain activity, to classical DCM and a purely behavioral model, we also investigated whether bDCM parameters encode additional information about the task that is not comprised in DCM or the Rescorla-Wagner model. For this, we followed the idea of generative embedding (Brodersen, Haiss, et al., 2011; Brodersen, Schofield, et al., 2011) and tested whether we could separate the horizontal and vertical runs of our experiments based on the parameter estimates of our models, which was not possible in previous analyses (Steinkamp et al., 2020).

## Methods

### Participants

Data were collected from 29 participants (15 female, 21-39 years old, M=25, SD=3) with normal or corrected-to-normal vision (all right-handed, Edinburgh handedness Inventory (Oldfield, 1971), M=0.86, SD=0.14), who provided written informed consent to participate in the study. One participant had to be excluded subsequently because of noncompliance. Another participant was excluded due to excessive head-movement (predefined criteria translation > 3mm, rotation > 3°). Furthermore, we could not extract the time-series for the left-TPJ VOI in one participant. Therefore, the final sample included 26 participants. The ethics board of the German Psychological Association had approved the study. Volunteers were paid 15€ per hour for their participation. The dataset has been used in a previous study (see Steinkamp et al., 2020).

### Task

Participants performed a spatial cueing task while lying in a 3T Trio (Siemens, Erlangen) MRI scanner. Stimuli were displayed on a screen behind the scanner bore, which could be seen via a mirror (mirror to display distance: 245 cm) that was mounted on a 32-channel head coil. The participants’ task was to report the orientation (horizontal/vertical) of a target Gabor patch (size 1° visual angle) by button presses of either the left or the right index finger while continuously fixating a diamond in the center of the screen (0.5° visual angle). A brightening of the central diamond (500 ms) indicated the beginning of a trial and was followed by a spatial cue after 1000 ms (brightening of one of the diamond’s edges for 200 ms) that indicated the location of the next target stimulus with 80 % probability. Participants were explicitly informed about the percentage of cue validity. The possible target locations were indicated by empty boxes (1° width) located to the left, right, top, and bottom of the fixation diamond (4° visual angle). After 400 ms or 600 ms, the target stimulus appeared for 250 ms at the cued location or in the box opposite to it. Distractor stimuli (constructed from two overlapping Gabor patches that were rotated by −45° and 45°, respectively) appeared simultaneously in the remaining three locations. Participants performed two runs of the spatial cueing-paradigm. In one run, targets and cues occurred along the vertical axis, in another along the horizontal axis (see Figure 1).

**Figure 1.**
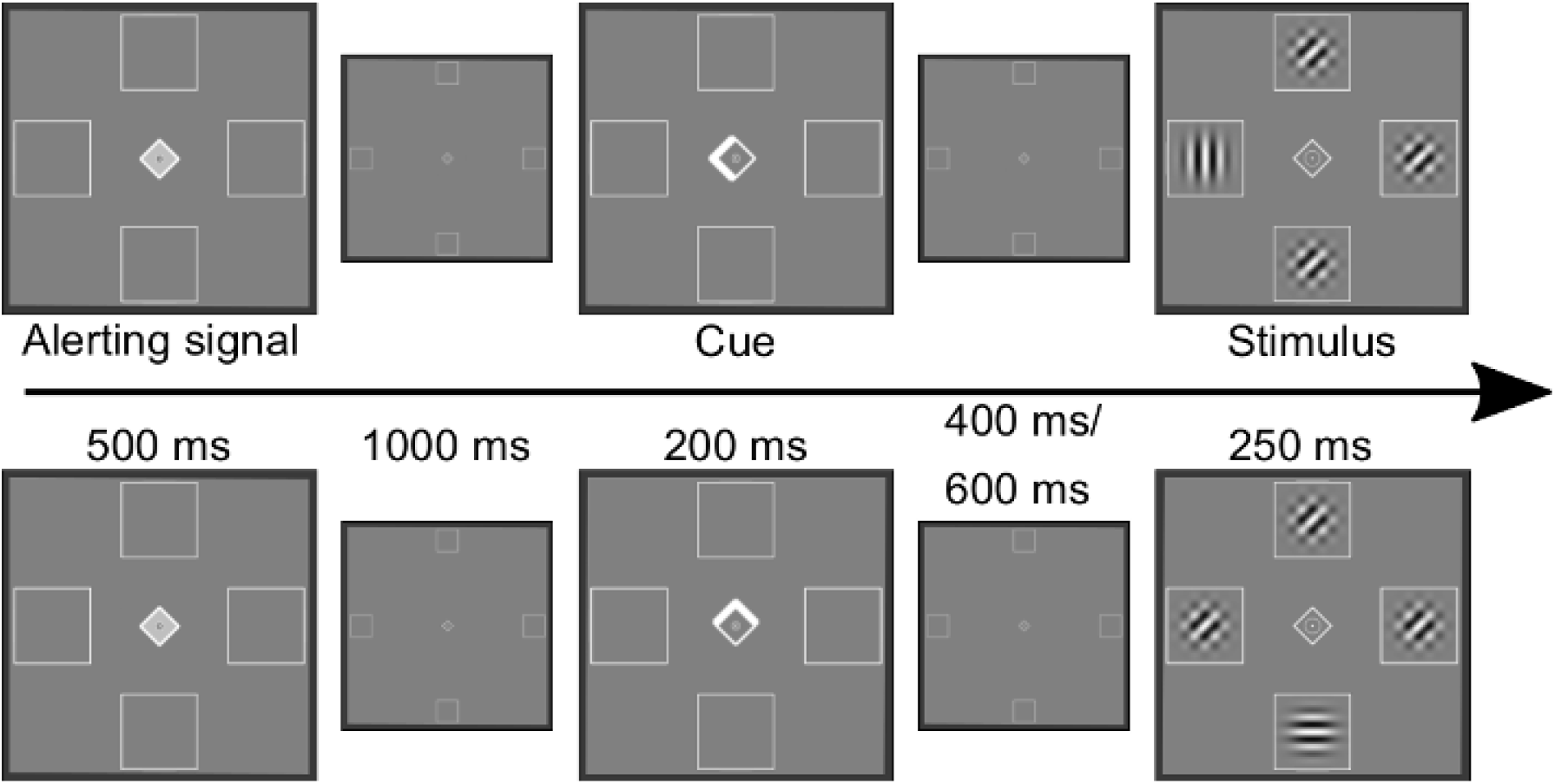
Illustration of the spatial cueing paradigm. In the upper row, a valid trial of the horizontal run is shown. The lower row depicts an example of an invalid trial in the vertical run. Reproduced from Steinkamp et al. (2020).

Each run consisted of five blocks of 40 trials (32 valid, 8 invalid). All possible combinations of target location, target orientation, and inter stimulus interval were presented in random order within each block. The time between the trials was drawn from the set of 2.0 s, 2.7 s, 3.2 s, 3.9s, or 4.5 s with equal probability. Between the blocks, there was a break of 10 to 13 s.

Run order (vertical or horizontal first) and the response mapping (left index finger for vertical orientations/right index finger for horizontal orientations or vice versa) were counterbalanced across participants. Before the experiment, participants performed a rapid detection task to train the mapping of stimulus-response associations. Here, targets appeared rapidly in the middle of the screen, and participants had to press the corresponding button as fast as possible. Immediate feedback and a running score of their accuracy were given. Additionally, there were 20 practice trials with feedback before each run of the main experiment.

Stimulus presentation and response collection were controlled using PsychoPy (version 1.85.3, Peirce, 2007, 2008; Peirce et al., 2019).

### Behavioral analysis

The mean reaction time was calculated for each participant and cueing condition and for each target location. Before calculating the mean reaction times, we preprocessed the data for each participant separately. First incorrect, missed, and outlier trials were removed. Outliers were defined as trials with reaction times below 0.2 s and reaction times greater than the 75^th^ percentile + 3 * Inter Quartile Range (IQR). The higher threshold for outlier exclusion was chosen to retain as many trials as possible in the analysis (removed trials, including errors, in the horizontal run: invalid M = 2.54, SD = 2.63; valid M = 6.62, SD = 5.91; in the vertical run: invalid M = 3.12, SD = 1.8; valid M = 6.0 SD = 3.94).

For the analysis of the “validity effect” (i.e., the slowing of reaction times in invalid as compared to valid trials), the data were pooled across the two runs (horizontal/vertical). The mean reaction times of the 2 x 4 (cueing x target location) factorial design were then analyzed in a repeated-measures ANOVA. The analysis was conducted in Python 3.7 using pingouin (version 0.3.3, Vallat, 2018).

### fMRI analyses

For each participant and each run, we collected 557 T2*-weighted images using an echo planar imaging (EPI) sequence (time of repetition (TR) 2.2 s; echo time (TE) 30 ms; flip angle 90°). Each recorded volume consisted of 36 transverse slices with a slice thickness of 3mm and a field of view of 200mm. The voxel size was 3.1 x 3.1 x 3.3 mm. The first 5 images were discarded to account for T1 equilibrium artifacts. Next to functional images, we also obtained an anatomical T1-weighted image for each participant, which was used in the preprocessing.

We preprocessed the fMRI data using fmriprep (version 1.1.1, Esteban et al., 2019), a robust and standardized pipeline, which applies slice-time correction, realignment, and normalization to MNI space. A detailed preprocessing report can be created automatically (see http://fmriprep.readthedocs.io/en/1.1.1/workflows.html), and has been included in the supplement.

Data was further spatially smoothed using an 8 x 8 x 8 mm FWHM Gaussian kernel. This step was done in Matlab 2018b (The MathWorks, Inc., Natick, Massachusetts, United States), using SPM12 (version 7771, Friston, 2007).

### fMRI - GLM

A classical GLM analysis was performed to identify activation peaks during attentional orientation and reorientation, which were later used to extract BOLD time-series data for the DCM analysis. The GLM analysis was conducted using SPM12. First-level models were created with four regressors of interest for each run, representing invalidly cued targets on the left (iL) and on the right (iR), as well as validly cued targets on the left (vL) and the right (vR) for the horizontal run, and invalidly and validly cued targets in the lower (iD, vD) and the upper (iU, vU) part of the screen in the vertical run.

To account for other physiological noise in the BOLD signal, we added the three rotation and three translation estimates of the rigid body transform, the average white matter signal, and the average cerebral spinal fluid (CSF) signal as nuisance regressors. We further included the squared time-series of the 8 regressors, the time-shifted time-series (t-1), as well as the square of the shifted time-series, resulting in a total of 32 nuisance regressors (Friston et al., 1996). We also applied a high pass filter at 128 s. For each run, four first-level contrasts were calculated: T-contrasts of valid and invalid trials versus baseline, an F-contrast of target onset versus baseline, which were used in the VOI analysis, and a differential contrast of invalid trials greater than valid trials. The latter contrast isolates brain regions involved in the attentional reorientation of attention.

At the group (second)-level, we investigated the differential contrast of invalid > valid trials using two planned one sample permutation t-tests against 0 using SnPM 13 (Nichols & Holmes, 2002), with default settings, 10000 permutations, and no additional variance smoothing. The cluster forming threshold was estimated during the processes with a predefined voxel-level cutoff of p < 0.001.

### Modeling Analysis

In the following, we will describe the modeling approaches used in our analysis, followed by a description of our model assessments and further analyses.

#### Rescorla-Wagner Model

We employed a variant of the Rescorla-Wagner model that we already used previously (Mengotti et al., 2017). While these studies were interested in the α parameter (the learning rate that describes how quickly participants adjust their internal assessment of the cue-validity), we applied this modeling approach to simulate reaction times in a trial-by-trial fashion. For parameter estimation, we defined new functions for the VBA (Variational Bayesian Analysis) toolbox (clone from master, in Jan. 2020, Daunizeau et al., 2014).

We used the following reinforcement learning formula as the evolution function, describing the hidden process governing the generation of reaction times.

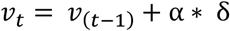

Where *δ* = *u_t_* – *v*_(*t*−1)_ describes the prediction error at trial t. The external input *u_t_* ∈ [0, 1] describes whether the cue at time t was either valid (1) or invalid (0), α is the learning rate and *v_t_* is the participant’s perceived cue validity after observation of trial t.

The observation function (i.e., the mapping from perceived cue validity to reaction times) was defined as:

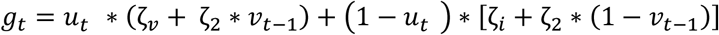

According to this formulation, the perceived cue-validity of the previous trial governs the responses, with different bias parameters for valid and invalid trials and a general scaling parameter of the predictions.

Table 1 depicts the Gaussian priors used in our estimation:

**Table 1:** Overview of parameters and prior values of the Rescorla-Wagner Model.

| Parameters | $\mu$ | $\sigma$ | |
| --- | --- | --- | --- |
| $\alpha$ | 0.5 | 0.5 | To ensure $0 < \alpha \leq 1$, $\alpha$ was logit and inverse logit transformed during parameter updating |
| $\zeta_v$ | 0 | 1 | |
| $\zeta_i$ | 0 | 1 | |
| $\zeta_2$ | 0 | 1 | |
| $v_0$ | 0.5 | 1 | Initial state of v |

### Behavioral DCM

In the following, we will provide a short overview of key-concepts of dynamic causal modeling (DCM). For a full derivation and detailed description of DCM, see (Friston et al., 2003; Rigoux & Daunizeau, 2015; Stephan et al., 2008). DCM is a fully-Bayesian approach to create a generative model of brain dynamics and infer effective connectivity between selected brain regions. In principle, DCM describes how experimental variations (described by the input u) drive the neural activity (x, the hidden states) in brain regions of interest in a dynamical system. The evolution function (*ẋ* = f(*x, u*)) describes the temporal dynamics of the hidden states (*ẋ*) and how they are influenced by external inputs (u). In DCM for fMRI, the evolution function f is typically described as:

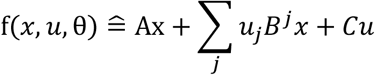

Where j corresponds to the number of inputs and i to the number of brain regions. The neural evolution parameters in θ correspond to the entries in A (fixed connectivity between brain regions), *B^j^* (modulation of connection strength by input j), C (direct effects of inputs). Hemodynamic states z (dependent on the neural states x) are then gated through an observation function:

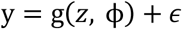

This function captures BOLD signal variations based on the hemodynamic states (z) and the hidden neural activity (x), with hemodynamic parameters ϕ. This mapping allows to observe and infer the hidden neural dynamics via the BOLD signal.

BDCM augments the described formulation of DCM by an additional evolution (h(*x, u*, ψ)) and observation functions (*g_r_*(*r*) + *ϵ_r_*) to map the hidden neural dynamics to behavioral responses. The evolution function h of the new “behavioral” state follows the same rationale as the function f in the DCM formulation:

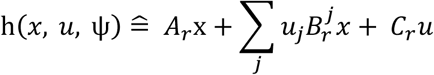

Here the parameter vector ψ describes the linear (*A_r_*) components of the behavioral state, as well as the direct (*C_r_*) and modulatory 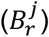 influences of experimental manipulations. *A_r_* is an analogy of the weight vector in a regression model. In the original paper, the neural states were mapped to binary behavioral observations (button press absent or present) via a sigmoidal function:

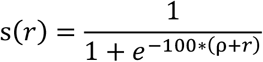

Here, ρ is an unknown bias term and r is the response or decision state. In our study, we slightly adjusted the sigmoid mapping, by changing the scale on which it operates. As we are not expecting reaction times slower than 3 s, we used this as an upper bound:

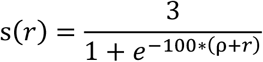

### Regions

As in our previous study (Steinkamp et al., 2020), we included bilateral IPS and FEF in our DCM model, which correspond to the central nodes of the dorsal fronto-parietal attention network (Vossel, Geng, et al., 2014). Additionally, as part of the ventral attention network, we included the TPJ bilaterally. As additional inclusions (e.g., the inferior/middle frontal gyrus) would have led to increasing model complexity and computational resources (and time), we did not include other brain regions, which may also play a role in attentional reorienting.

Based on our assumptions about the dorsal and ventral attention network’s interplay, we created three automatic meta-analysis using Neurosynth (https://www.neurosynth.org/, Yarkoni et al., 2011), to define the seed coordinates for the subsequent VOI analysis (see Table 2). Our regions of interest were bilateral IPS (search term: “intraparietal sulcus”), bilateral FEF (search term: “frontal eye”), and bilateral TPJ (search term: “tpj”). We downloaded the corresponding association maps (associations, p<0.01 FDR corrected) and identified the seed location as the peak voxel in the cluster of interest, using the Anatomy toolbox (v2, Eickhoff et al., 2005). In all three maps, the two largest clusters encompassed our regions of interest in either the left or right hemispheres.

**Table 2:**
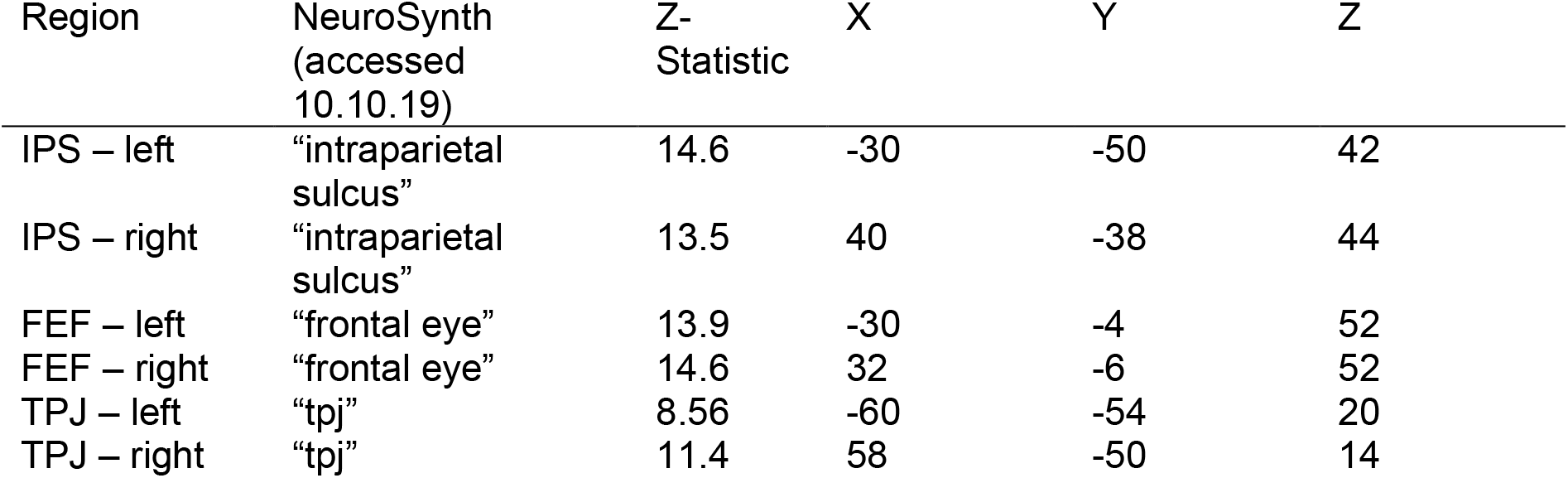
Regions and search-terms for automated Neurosynth meta-analyses.

In each run, we used the participant level t-maps (thresholded at p < 0.1 uncorrected) to search for individual local maxima in a 12 mm sphere around the seed coordinates. The first principle component of BOLD time-courses in a 9 mm VOI around the participant’s maximum was extracted and adjusted based on the F-contrast defined in the first-level analysis. Task-related activity for the IPS and FEF VOIs was defined by the contrast of valid trials against baseline and for TPJ by the contrast of invalid trials against the baseline.

### Preprocessing

We preprocessed the BOLD signal by detrending the signal in each VOI (*spm_detrend*) and scaling the BOLD amplitude across VOIs to a maximum value of 4 (see *spm_dcm_estimate*). Behavioral data were extracted from the event data, and as in the previous analyses, error trials and trials with missed responses, as well as RTs fulfilling the outlier criterion (RT < 0.2s and RT > 3 * IQR + UQ), were excluded.

BOLD data were resampled from a TR of 2.2 s to a sampling rate of 1.1 s (by interspersing “NaN” values). The behavioral observations were set to occur at the corresponding target onset, which was also downsampled to a resolution of 1.1 s. No resampling of BOLD data was performed for the classical DCM analysis. As the Rescorla-Wagner model represents trial-by-trial dynamics, the corresponding preprocessed reaction times were used, excluding error and missed trials.

For our modeling, we assumed homogenous HRF dynamics across the six regions, fixing the initial states of the model to 0 and estimating the shape of the observation noise hyper-prior distributions. For this, we assumed that we would be able to explain 10 – 90% of the variance in both the BOLD and the reaction time data. The prior distributions over the other parameters were set to the defaults of the VBA toolbox. We used the same hyperpriors for the explained variance of the BOLD signal in the classical DCM analysis and the Rescorla-Wagner model’s behavioral responses.

To define the inputs into the DCM models, we created separate SPM-design matrices that were only used to define the input streams. Stream one was defined as the driving input to all six regions, containing an impulse every time a target stimulus appeared (irrespective of the cueing condition or target location). The second stream was used purely for the modulatory effects, containing an impulse only in invalidly cued targets. The input streams were extracted from the SPM design matrix and were centered before entering the model inversion (*spm_detrend*). As mentioned above, the Rescorla-Wagner model is modeling trial-by-trial variations (rather than continuous time), so the input to this model was a vector consisting of ones and zeros, indicating whether the current trial is invalid or valid.

### Model definition

We used the same general model structure for behavioral and classical DCM analysis. As in our previous publication, we used IPS, FEF, and TPJ as our brain regions of interest. For our analysis, we inverted a single model. The fixed connectivity structures of our model (i.e., the A-matrix) had full connections in each hemisphere and connections between homologous regions (Figure 2). As we did not include visual areas in our modeling approach, all six regions received driving input (C-matrix). For bidirectional intra- and interhemispheric modulatory connections (B-matrix), we considered the IPS and TPJ. Connections in both hemispheres to the FEF were unidirectional, assuming that there were no feedback modulations from FEF to the other brain regions. In the case of bDCM, we also considered all six nodes as output regions.

**Figure 2:**
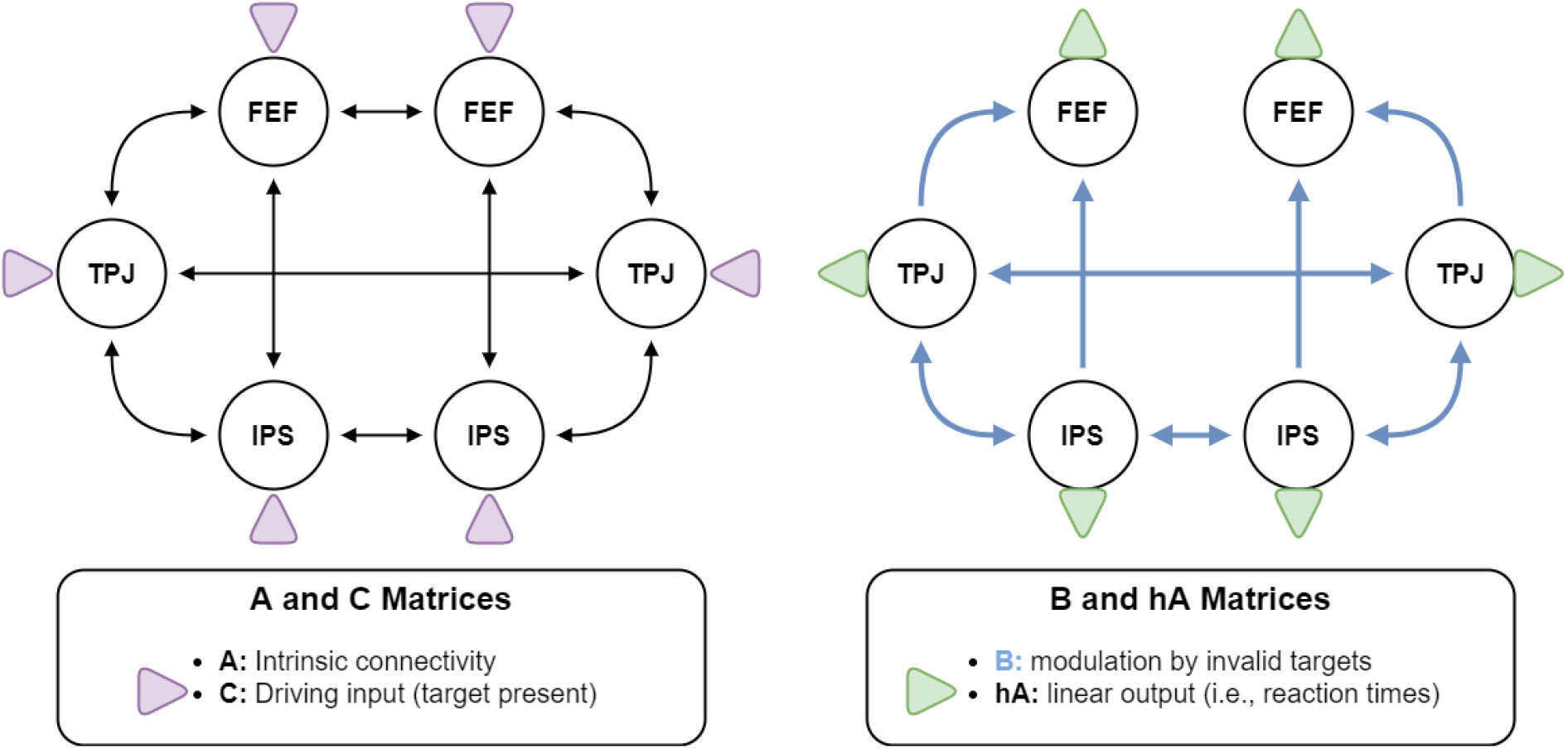
Basic structure of the DCM model. Regions were fully inter-connected in each hemisphere, and homologous regions were connected. All regions received driving input. We assumed that all connections between the regions were modulated by invalid trials, except for feedback and interhemispheric connections from FEF.

### Model evaluation of the Rescorla-Wagner model, classical DCM, and bDCM

As all three models are based on different underlying data at different times scales (i.e., reaction times only, BOLD time-series only, or both), we only compared the models based on their outputs, applying classical goodness of fit-statistics. The R^2^-score,

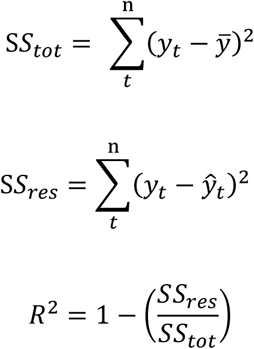

where *y_t_* describes the datapoint at *t*, with *n* timepoints in total. The average of y is defined as 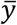, and *ŷ* are predicted values. Similarly, we also calculated the mean absolute error (MAE)

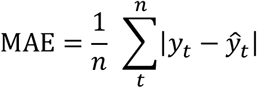

Here, we estimated for each subject whether the fit-statistics were different from random for each of the model outputs by permutation testing. The predicted values *ŷ* were shuffled 10000 times (without replacement), and the two statistics were recalculated. The permutation p-value for the models is then reported as the proportion of fits greater than the model’s R^2^–score (smaller in case of MAE) plus one divided by the number of permutations plus one (Ojala & Garriga, 2010). At the group level, we report the proportion of significant models, based on a permutation p-value < 0.05.

To compare the model performance on reaction times, we used (Bayesian-) paired t-tests to test for differences between the fit-statistics (R^2^-score and mean absolute error (MAE)), separately for the two runs. We then investigated how well bDCM and the Rescorla-Wagner model simulate the underlying reaction time distributions. This was achieved by calculating a two-sample Kolmogorov-Smirnov test between the model-derived reaction times of the Rescorla-Wagner model or bDCM and the measured reaction times. Finally, paired t-tests were used on the distance between the distributions (as determined by the KS-test) to test which simulation followed the measured data more closely (i.e., had a smaller distance at the group level).

We applied a mixed-effects linear model for each error term to compare differences in performance to the BOLD data fit between classical DCM and bDCM. The mixed-effects model followed the following formula, where “Score” either depicts the mean absolute error or the R2-score:

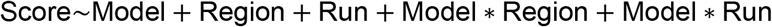

and Model has the two factors “DCM” and “bDCM”, “Run” describes either the horizontal or vertical run, and “Region” indicates the “Score” for either VOI. Each model also contained a random intercept for each participant.

### Generative Embedding

We also investigated whether bDCM parameters encode additional information, which is not contained in classical DCM or in the Rescorla-Wagner model. Therefore, we tested whether the parameters derived from our computational models could be used to separate the horizontal and vertical run. Note that our previous work using classical DCM did not provide any evidence for distinct behavioral or neural processes (Steinkamp et al., 2020).

To test if a separation is possible and which sets of parameters encode the necessary information, we ran several experiments using different feature sets. Our feature sets for DCM and bDCM were: “AB”, the sum of the “A” and “B” matrices (19 parameters); “AB + C”, extending the feature set with the “C” matrix parameters (25 parameters); in case of bDCM we additionally tested the feature sets “AB + hA”, where the “hA” matrix was also included in the set (25 parameters); “AB + C + hA”, including all connectivity parameters of the bDCM (31 parameters); “C + hA”, using only the input and outputs connections of bDCM. We then also tested whether we can find any information in the parameters of the Rescorla-Wagner model (see Table 2, “RW”, 4 parameters), and whether parameters of DCM and the Rescorla-Wagner model in combination have a beneficial effect (“AB + C + RW”, 29 parameters). To be sure that there was no additional information in the BOLD time-series, we also included a feature set based on the correlation of the time-series of the six VOIs in our experiments (“Correlation”, 15 parameters) and of the time-lagged correlation with the time-series (“Correlation + Correlationt-1”, 51 parameters). Correlations were established using nilearn’s *ConnectivityMeasure* (Abraham et al., 2014), with no variance scaling.

As we have a minimal sample size for classification approaches (26 participants, 52 instances), we applied an elaborate cross-validation procedure for our results. We applied 5-fold cross-validation to obtain an estimate of the accuracy of each feature set, whose significance was further evaluated by permutation testing (1000 iterations, scikit-learn’s, *permutation_test_score*; Pedregosa et al. (2011)). To keep more independence between the train and test-sets, we ensured that a given participant’s instances were not distributed across training and test-sets. Due to the small sample size and high variance between participants, we repeated the cross-validation procedure 20 times, shuffling the participants order, and creating different train-test splits for each iteration.

To account for a certain amount of algorithmic variance, we performed the procedure above using two classifiers, a logistic regression, and a linear support vector machine (scikit-learn, using default parameters). Input data were normalized using robust-scaling (scikit-learn’s *RobustScaler*).

### Lesion Analysis

We also applied lesion analysis to the bDCM model, as described in Rigoux and Daunizeau (2015). Here, the afferent connections towards a single brain region were reduced to 0 to simulate the absence of this region (i.e., to create an artificial lesion). The simulated data from such a lesioned model can be used to better understand behavioral changes after damage to certain brain regions.

To alleviate the problem of numerical instabilities, which resulted in values resulting in infinity or minus infinity for the hidden states, we changed the posterior self-inhibitory connection in the DCM model to log(1) (to ensure inhibition, i.e., negativity, self-connections in DCM are exponentiated before subtraction from the diagonal of the A matrix). While increasing the self-inhibition solved the problem of instabilities, it significantly impacted the fit-statistics of the non-lesioned model. Therefore, we only provide a qualitative description of the lesion analysis. In the end, we simulated data for all 26 participants for each lesion and several levels of lesion extent. This means, rather than switching off the afferent connections (i.e., the inputs to the region) altogether, we also simulated data for connections that had 95 %, 75 %, 50 %, 25 %, 5%, and 0 % of the original incoming strength.

Since a few models still were numerically unstable, we then cleaned the simulated data by removing datasets on a per lesion basis where the variance after the 20^th^ trial was close to 0 (i.e., the simulated reaction times flatlined at the maximum/minimum of the sigmoid function) and which returned non-values. In the qualitative analysis, we compared the group-mean validity effects for different lesions, extents, and target sides, removing participants who had an absolute validity difference greater than 2 secs.

## Results

### Behavior

To test for reaction time effects of cueing (valid or invalid) and target-side (left, right, down, up), as well as their interactions, we applied a 2 x 4 repeated measures ANOVA (see also Figure 3). There was no significant effect of target-side (F(1.965, 49.125) = 0.1, p = 0.902, 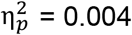, ε = 0.655). However, a significant main effect of cueing (F(1, 25) = 26.647, p < 0.001, 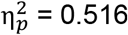, ε = 1.0) and a weak significant interaction between target-side and cueing (F(2.88, 72) = 2.866, p = 0.045, 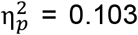, ε = 0.96) were observed. All reported p-values were Greenhouse-Geisser corrected to account for a lack of sphericity.

**Figure 3:**
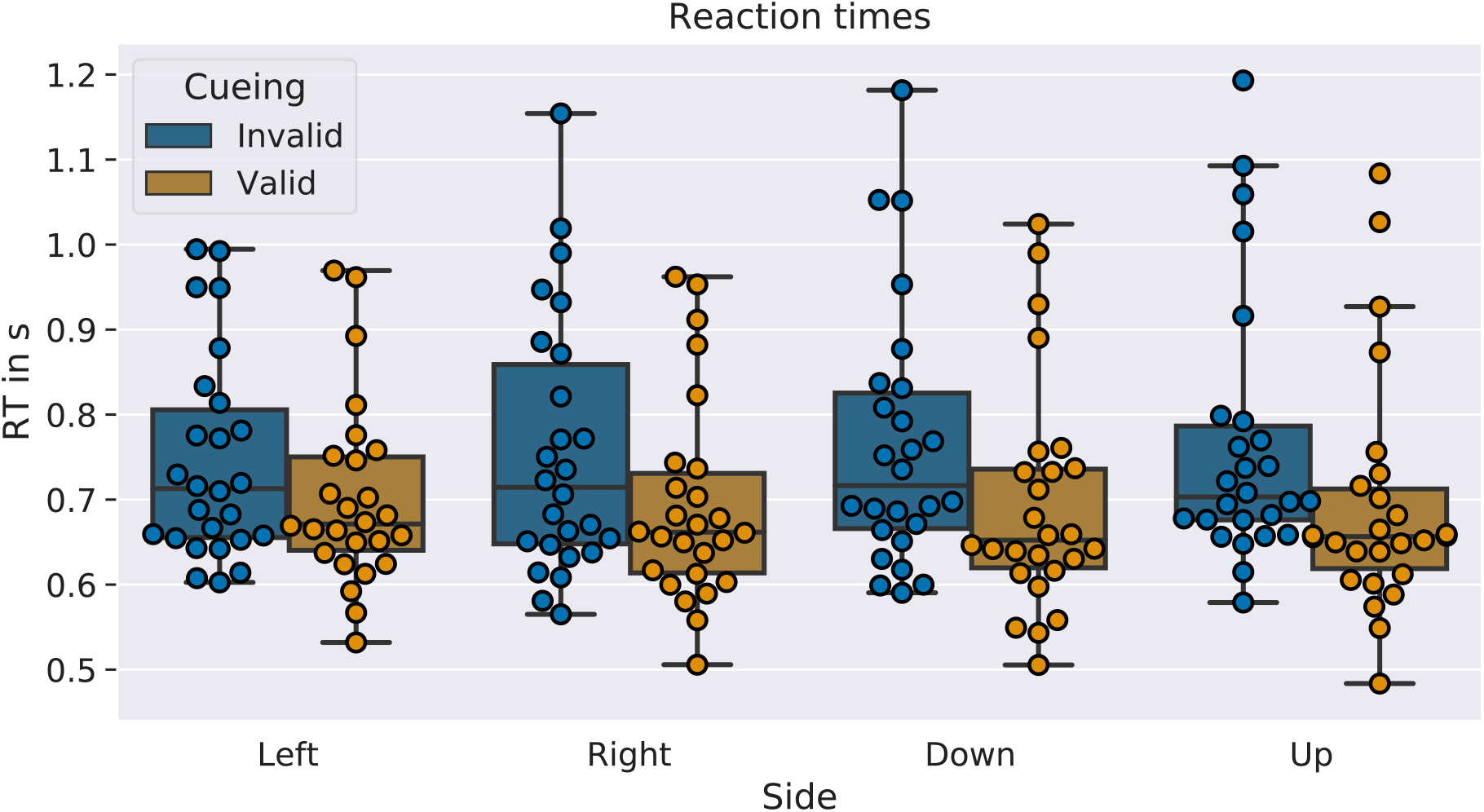
Box- and swarm plots of mean-reaction time data for each participant in the 8 conditions. The boxes indicate the inter-quartile range (IQR), the line in the middle the median reaction time, whiskers are extended to include the lower and upper quartiles plus 3 times the IQR. Loose points indicate outliers. The ANOVA’s results are readily visible, as there are longer reaction times in invalid trials but no apparent effects between the different target-positions.

### FMRI GLM

The contrasts of invalid versus valid trials isolating reorienting-related activity for the two runs are reported in Figure 4 (group t-maps are provided on neurovault:

**Figure 4:**
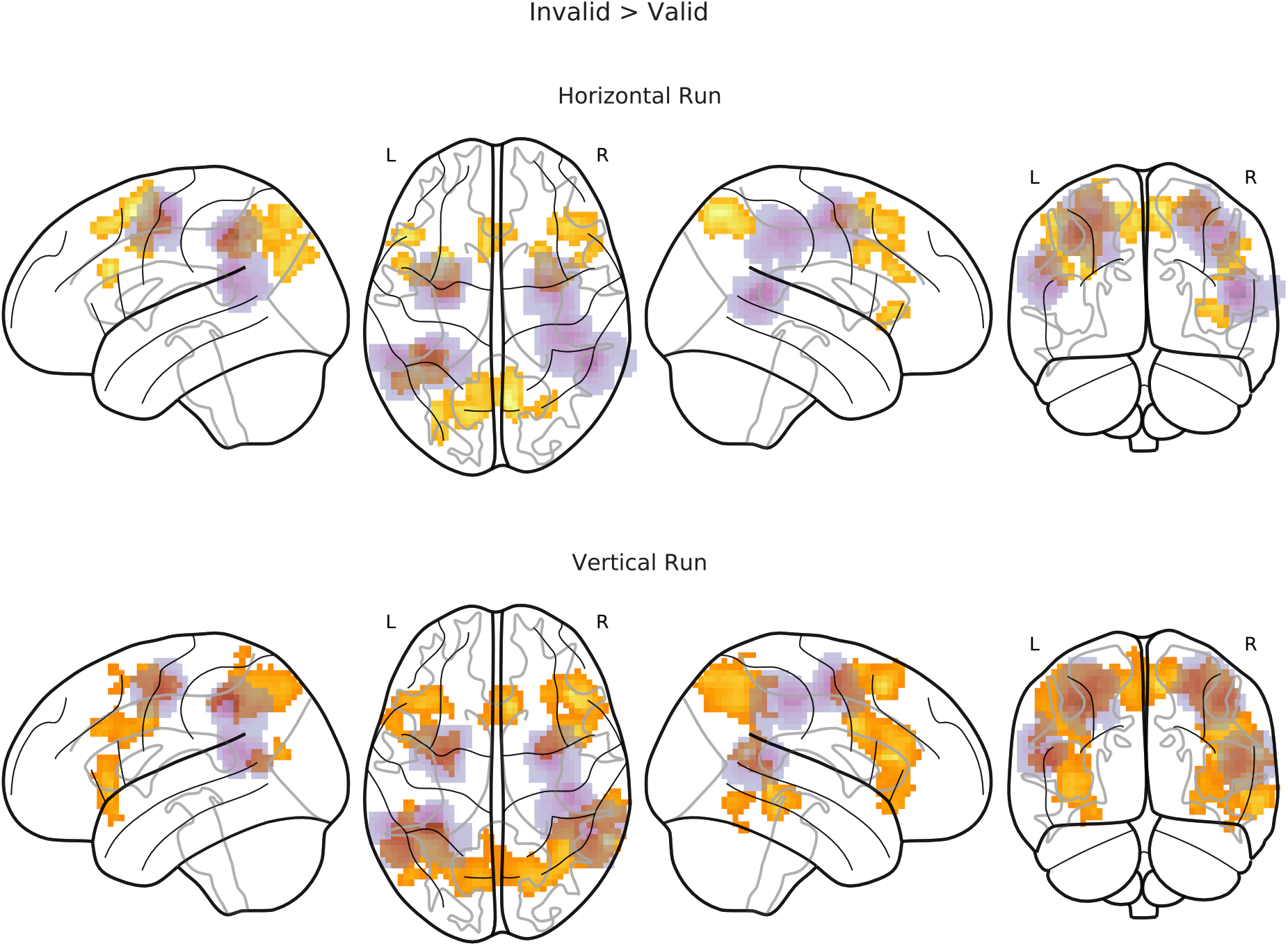
Non-parametric T-maps contrasting invalid > valid trials for the two runs (p < 0.05 FWEc). The purple overlay indicates the regions where the 9mm VOIs for the bDCM analysis were extracted (sum of the participant’s masks).

NV_LINK, (Gorgolewski et al., 2015), the corresponding tables reporting global and local maxima for the different clusters are in supplement S2). We performed a one-sample permutation t-test on the first-level contrast images (invalid > valid), with a predefined cluster-forming threshold of p < 0.001, the results are reported family-wise error corrected at p < 0.05 (cluster threshold horizontal k = 58 voxels, vertical k = 70 voxels). In both maps, we found areas classically associated with the dorsal and ventral attention networks. For example, in both runs, we observed significant activation in bilateral intraparietal sulci and frontal-eye fields. Activations of the ventral attention networks were less robust. For the horizontal run, for example, the invalid versus valid contrast revealed an involvement of the middle frontal gyrus predominantly in the right hemisphere and no significant activation close to the seed regions for the temporoparietal junction at the given threshold. However, the temporoparietal junction was significantly activated in the vertical run.

### Model Fit

#### Reaction Time Data

We calculated the mean absolute error (MAE) and the R^2^ statistic for the Rescorla-Wagner and bDCM models and assessed their significance on a per-subject level by calculating permutation tests. In the horizontal run, the bDCM (mean absolute error, M = 0.091, SD = 0.033, percent sig = 96.2 %; R^2^ score, M = 0.117, SD = 0.084, percent. sig = 100 %) performed well when compared to the Rescorla-Wagner model (mean absolute error, M = 0.094, SD = 0.033, percent sig. = 53.8 %; R^2^ score, M = 0.066, SD = 0.08, percent. sig = 69.2 %). The results of the vertical run yielded a very similar picture, where the differences between the bDCM (mean absolute error, M = 0.093, SD = 0.034, percent sig. = 100 %; R^2^ score, M = 0.158, SD = 0.143, percent. sig = 100 %) and the Rescorla-Wagner model (mean absolute error, M = 0.096, SD = 0.038, percent sig. = 80.8 %; R^2^ score, M = 0.094, SD = 0.141, percent. sig = 88.5 %) were even more pronounced (see Figure 5). The paired t-tests (Table 3) confirmed this pattern and revealed a better fit for the vertical run. BDCM had a lower error and greater fit than the Rescorla-Wagner model.

**Figure 5:**
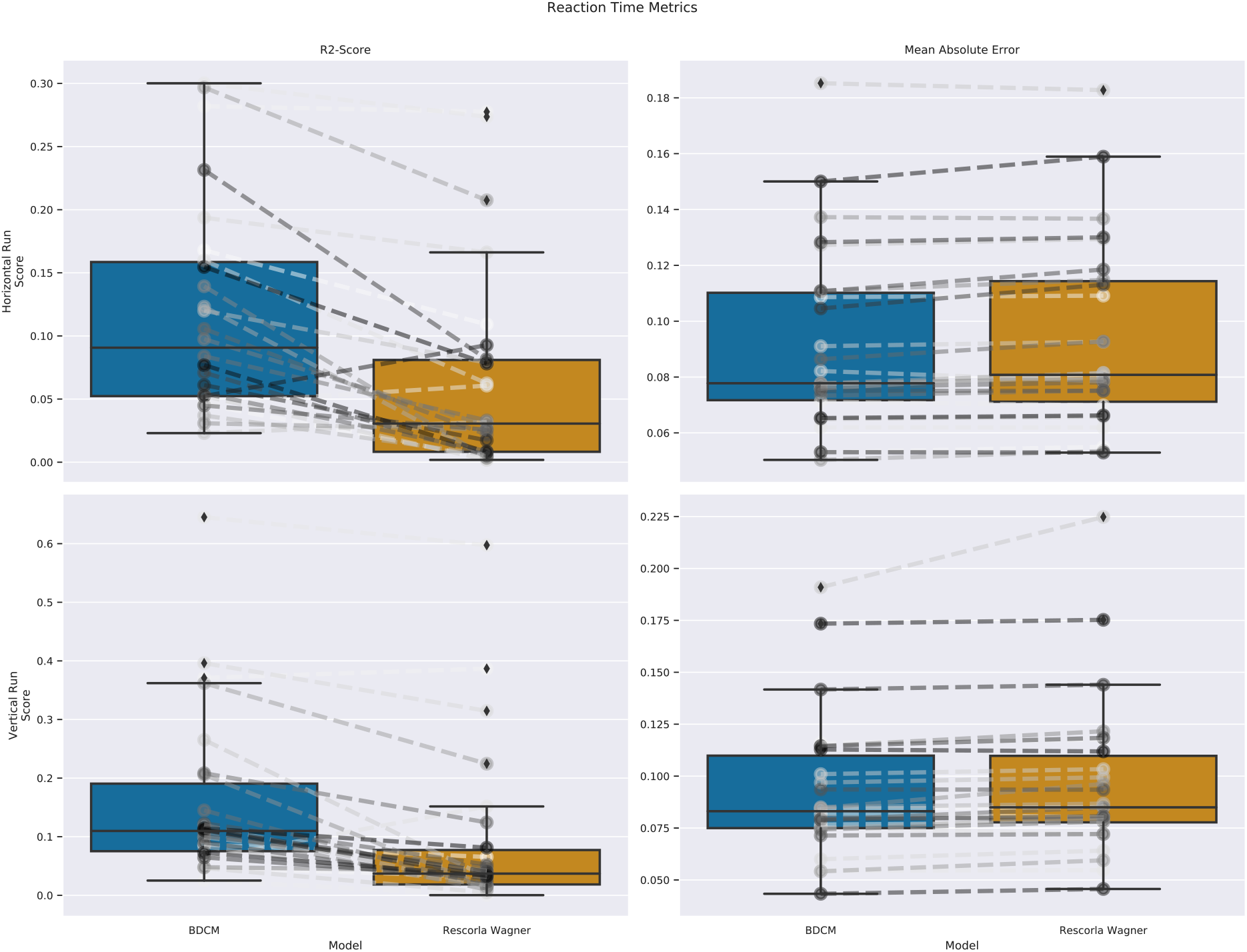
Boxplots comparing the different fit-statistics across models in the horizontal run. Please note that for the R^2^-score a higher value is better, while the opposite is true for the mean absolute error. The dashed lines between the boxplots indicate individual participants.

**Table 3:** Paired t-tests between the fit-statistics of bDCM and Rescorla-Wagner models. The differences between the fit-statistics favor the bDCM model.

| Run | Error | T (df 25) | p-val | CI95% | cohen-d | BF10 |
| --- | --- | --- | --- | --- | --- | --- |
| Horizontal | MAE | -3.557 | 0.002 | -0.0, -0.0 | 0.066 | 23.692 |
| Horizontal | $R^2$ | 5.803 | < 0.001 | 0.03, 0.07 | 0.625 | 4.249.729 |
| Vertical | MAE | -2.644 | 0.014 | -0.01, -0.0 | 0.098 | 3.571 |
| Vertical | $R^2$ | 5.204 | < 0.001 | 0.04, 0.09 | 0.456 | 1.048.508 |

Furthermore, we evaluated how well the reaction time distributions of the two models’ simulations matched the real reaction time distribution (Figure 6). We calculated the distance between the distributions of measured and simulated reaction times for each run, participant, and cueing-condition using the Kolmogorov-Smirnov test. We then performed paired t-tests in order to ascertain which simulation better matches the original distribution. In all cases, the bDCM simulation provided a better match (horizontal run, valid cueing, t(25) = −10.155, p < 0.001, Cohen’s d = 2.432, BF_10_ = 4.39 * 10^10^; horizontal run, invalid cueing, t(25) = −8.121, p < 0.001, Cohen’s d = 2.392, BF_10_ = 7.51 * 10^8^; vertical run, valid cueing, t(25) = −7.619, p < 0.001, Cohen’s d = 2.049, BF_10_ = 2.56 * 10^8^; vertical run, invalid cueing, t(25) = −9.931, p < 0.001, Cohen’s d = 2.475, BF_10_ = 2.87 * 10^10^). Based on visual inspection of the differences in reaction time distributions, deviations were especially pronounced at the extreme ends of the distribution.

**Figure 6:**
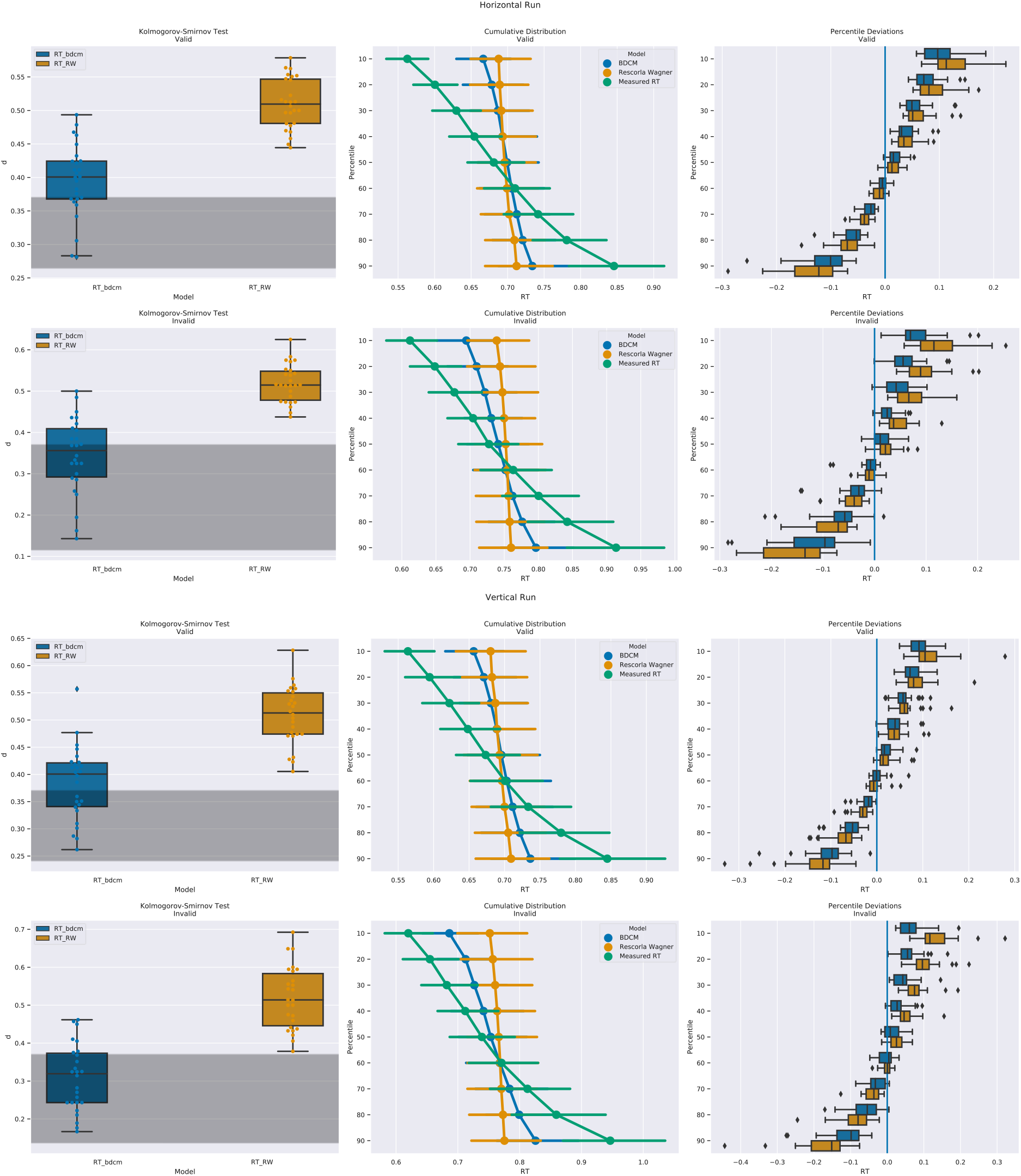
Left column, Kolmogorov-Smirnov distance between the simulated responses between the measured reaction times and bDCM or the Rescorla-Wagner model. The grey shading indicates non-significant differences. BDCM has, in general, a lower distance to the original distribution, and in many cases, the tests were non-significant. Both models also appeared to be better in the prediction of the invalid reaction time distribution. Middle Column: Cumulative reaction time distribution represented by the deciles of each model. The measured reaction times (green) have a more extensive spread than the reaction times from bDCM (blue) and the Rescorla-Wagner model (orange). Right column: The paired difference between the deciles of the Rescorla-Wagner model and bDCM. Differences are especially large in more extreme deciles.

### BOLD Data

In a further step, we investigated whether DCM and bDCM were comparable in their fit to the measured BOLD data. For this, we calculated the fit-statistics (mean absolute error and R^2^-score) for the two modeling approaches and the two runs. Since this yields fit-statistics for each brain region, we calculated two linear mixed-effects models (MLM) with participant as a random factor to test for a main-effect or interaction effects of model and fit-statistic, as well as the main effect of run and interactions between model and run (Figure 7). The results of the mixed-effects models are summarized in Table 4. Importantly, we did not find a significant main effect of model. However, there were significant main effects of brain region, these effects did not interact with the choice of model, indicating that the two models performed similarly. A similar conclusion can be drawn when looking at the different runs. Again, there were no significant main effects nor interactions of the factor model.

**Figure 7:**
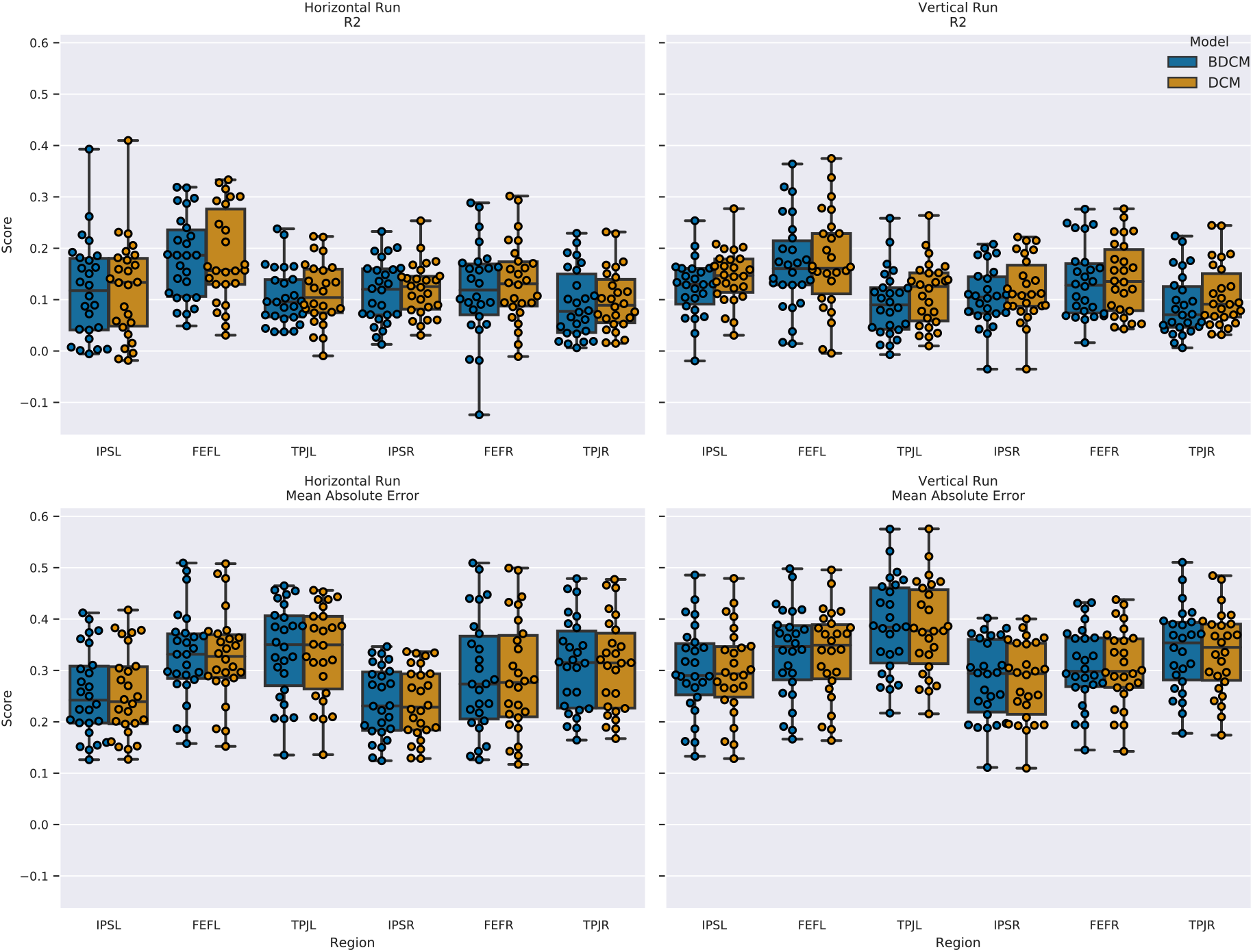
Box plots for the different fit-statistics. Left column shows the data for the horizontal run, the right data for the vertical run. The upper row indicates the R^2^ scores and the lower row the mean absolute error.

**Table 4:** Results of the MLM analysis for BOLD fit-statistics. The table is split for mean absolute error and R^2^ score.

| | $R^2$ Score | | | | Mean Absolute Error | | | |
| --- | --- | --- | --- | --- | --- | --- | --- | --- |
|  | Coef. | Std.Err. | z | P>\|z\| | Coef. | Std.Err. | z | P>\|z\| |
| Intercept | 0.176 | 0.012 | 14.986 | < 0.001 | 0.317 | 0.014 | 22.720 | < 0.001 |
| DCM | 0 | 0.013 | -0.017 | 0.986 | 0 | 0.016 | -0.021 | 0.983 |
| FEF Right | -0.048 | 0.012 | -3.879 | < 0.001 | -0.033 | 0.015 | -2.189 | 0.029 |
| IPS Left | -0.053 | 0.012 | -4.268 | < 0.001 | -0.054 | 0.015 | -3.574 | < 0.001 |
| IPS Right | -0.063 | 0.012 | -5.109 | < 0.001 | -0.072 | 0.015 | -4.764 | < 0.001 |
| TPJ Left | -0.075 | 0.012 | -6.074 | < 0.001 | 0.03 | 0.015 | 1.946 | 0.052 |
| TPJ Right | -0.083 | 0.012 | -6.727 | < 0.001 | -0.007 | 0.015 | -0.436 | 0.663 |
| Vertical Run | -0.003 | 0.007 | -0.411 | 0.681 | 0.031 | 0.009 | 3.593 | < 0.001 |
| DCM * FEF Right | 0.007 | 0.017 | 0.428 | 0.669 | 0 | 0.021 | 0.002 | 0.999 |
| DCM * IPS Left | 0.008 | 0.017 | 0.483 | 0.629 | -0.001 | 0.021 | -0.033 | 0.973 |
| DCM * IPS Right | 0.005 | 0.017 | 0.301 | 0.763 | -0.001 | 0.021 | -0.044 | 0.965 |
| DCM * TPJ Left | 0.01 | 0.017 | 0.552 | 0.581 | -0.002 | 0.021 | -0.087 | 0.93 |
| DCM * TPJ Right | 0.009 | 0.017 | 0.501 | 0.617 | -0.001 | 0.021 | -0.059 | 0.953 |
| Vertical Run * DCM | 0.008 | 0.01 | 0.801 | 0.423 | -0.002 | 0.012 | -0.137 | 0.891 |

### Generative Embedding

In this analysis, we tested how well the different models could separate the vertical and horizontal runs from each other using two different kinds of machine learning algorithms. Due to the high variability in our data (a very low number of training instances, difficult task), we decided to shuffle the assignment of train-test folds so that the cross-validation was performed multiple times. We assessed how often predictions of a classifier were significant (permutation P-Value < 0.05) and calculated the average predictive accuracy. The horizontal run could not be separated from the vertical run using combinations of features that were not derived from bDCM parameters (see Figure 8). This indicated that bDCM provided us with some very minute differences between the two runs, that were not included in other models or the BOLD signal.

**Figure 8:**
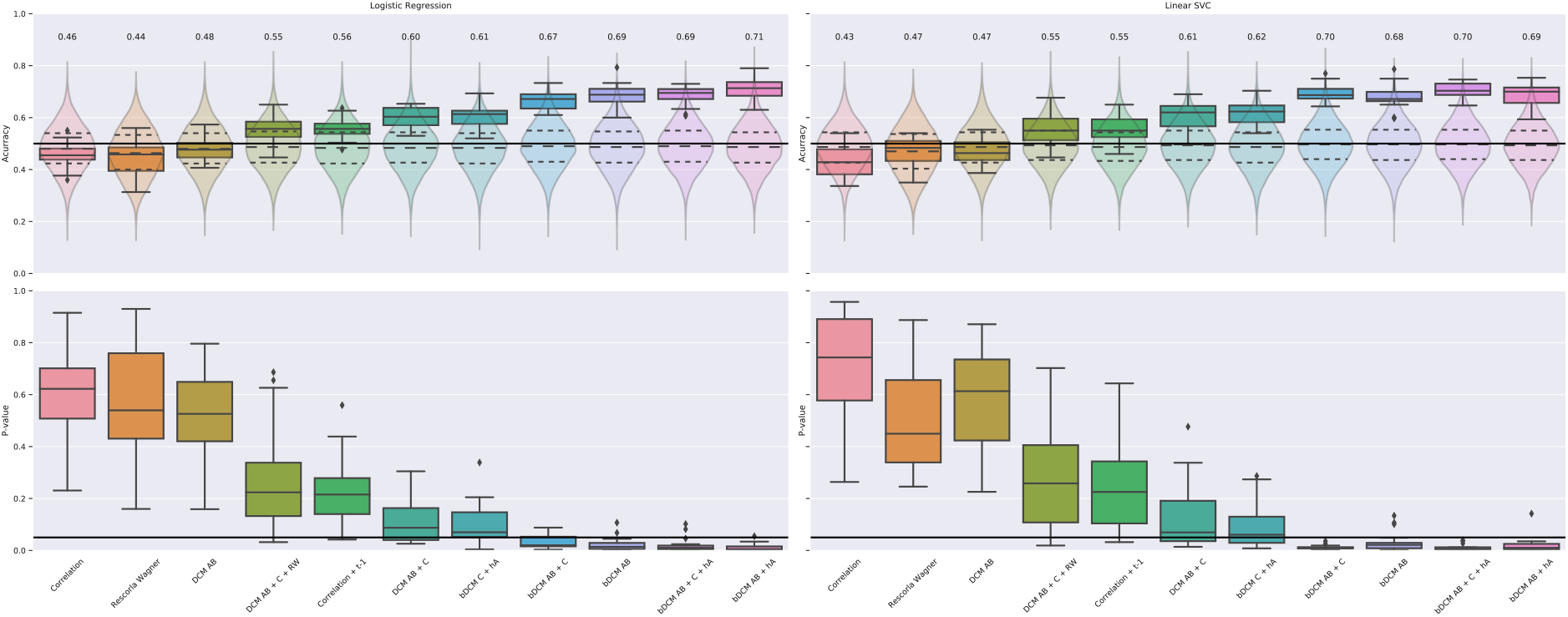
Classifier performance for the 20 shuffles of our dataset, sorted by average performance. In the upper row, the average performance and spread across iterations are shown, and transparent violin-plots indicate the distribution of permutation scores. In the lower panel, the distribution of permutation p-values is indicated, with the black line indicating the p < 0.05 cut-off.

### Lesion Analysis

Figure 9 depicts the results of the lesion analysis for the horizontal run (the vertical run can be found in the supplement). Constraining the self-connections in the model resulted in significant decreases in model fit of reaction time data in both the horizontal run (paired t-tests; MAE, t(25) = 5.054, p < 0.001, Cohen’s d = 0.197; R^2^-score, t(25) = −6.573, p < 0.001, Cohen’s d = 1.260) and the vertical run (paired t-tests; MAE, t(25) = 4.159, p < 0.001, Cohen’s d = 0.175; R^2^-score, t(25) = −7.185, p < 0.001, Cohen’s d = 0.898). Despite this note of caution, there were some interesting trends in the data. The effect of lesion extent on the validity effect seemed highly specific for the different network nodes with some lesions increasing and other lesion decreasing the validity effect, depending on the extend of the artificial lesion.

**Figure 9:**
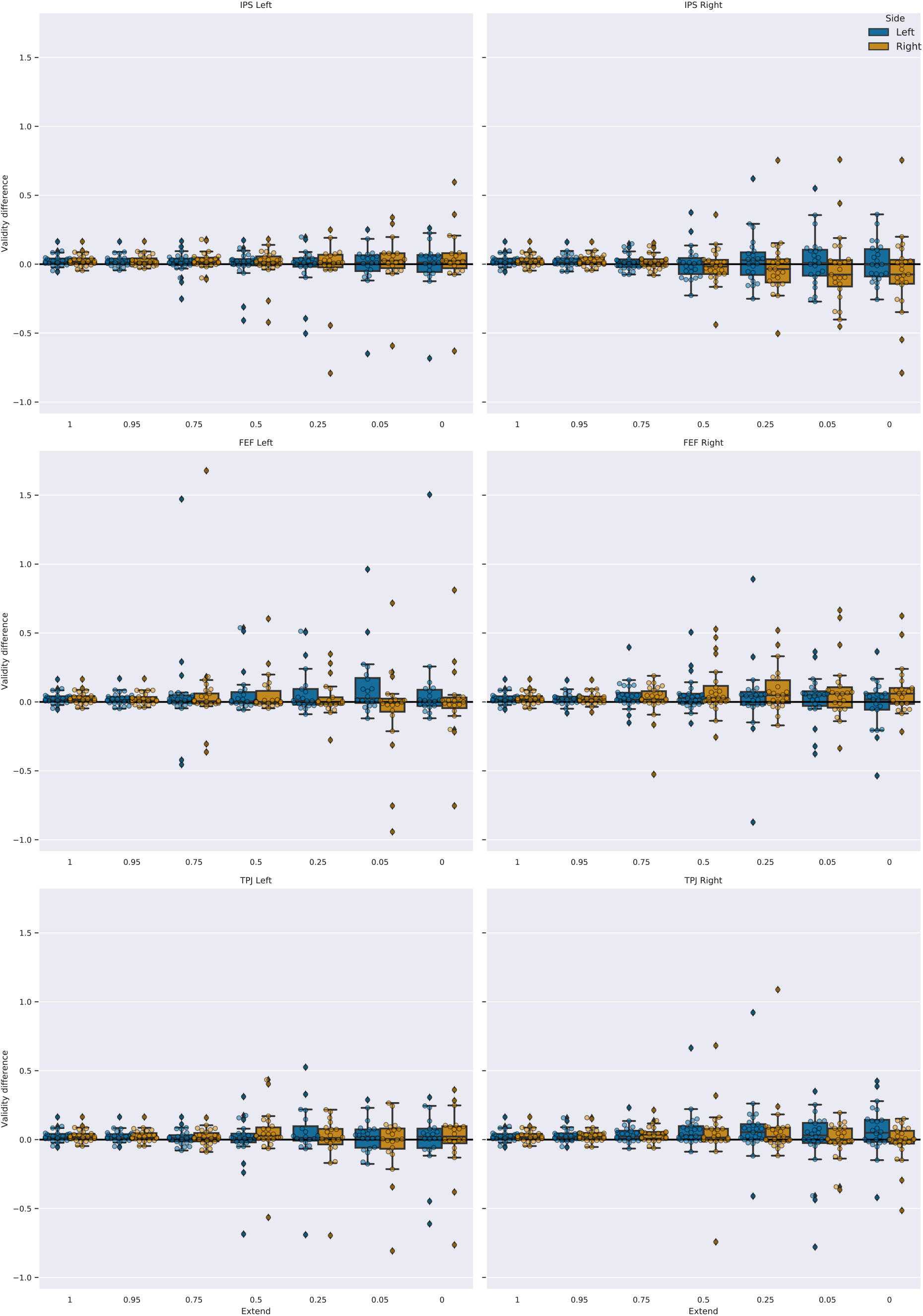
Validity effect for simulated reaction times for the horizontal run, after brain lesions of different extent. An extent of 1 means that no lesion occurred. An extent of 0 indicates that the region was fully disabled. Boxplots indicate the median of the data, the IQR, and the minimum and maximum values. Outliers exceed the 1.5 * IQR criterion.

From a computational anatomy perspective, lesioning the TPJ yielded plausible effects, with a gradual increase in the contralesional validity effect following larger lesions.

## Discussion

We applied bDCM (Rigoux & Daunizeau, 2015) to simultaneously model neural responses and reaction times in a spatial cueing task. We here demonstrated that bDCM could not only be applied to binary responses, but also to continuous read-outs (i.e., reaction times).

After reproducing previously published effects of cue validity at the behavioral and neural level, we modeled behavioral and functional imaging data in three different ways. BDCM, as a novel approach, was compared to both classical DCM and the behavioral Rescorla-Wagner model. As all three models serve different purposes and rely on different data on different timescales, we restricted the model comparison to the models’ outputs and fit-statistics.

Although the original paper on bDCM suggested that incorporating behavior also leads to an advantage in representing the BOLD response of bDCM over classical DCM, we did not find significant differences between both modeling approaches. The benefit of including behavioral measures might only be prevalent when BOLD recordings are noisier than behavioral recordings (Rigoux & Daunizeau, 2015).

Furthermore, we compared simulated reaction times of bDCM and our implementation of the Rescorla-Wagner model (Vossel, Mathys, et al., 2014). BDCM had in general a better fit to reaction time data (reflected in higher R^2^-score and lower error) and represented the distribution of reaction times more closely in both valid and invalid trials (reflected in significantly lower distances, which were calculated by the Kolmogorov-Smirnov test). Both the Rescorla-Wagner model and bDCM did not model the extreme ends of the reaction distributions well, however, bDCM deviated less from the measured data.

Note that this comparison was not performed to favor one model over the other. Instead, it was conducted to evaluate bDCM against the performance of a highly specialized, validated, and less complex model in a cueing task. Despite the superior fit of bDCM, the Rescorla-Wagner model performed extremely well, given the small number of parameters. Hence, if we penalized for model complexity, the Rescorla-Wagner model would probably be identified as the preferred model for reaction times. BDCM also incorporates the dynamics of the BOLD response and operates on a timescale of seconds, rather than trials. Thus, having only 8 parameters more than the classical DCM (69 parameters) seems to be an adequate increase in complexity. The resulting more detailed representation of reaction time distributions in bDCM might be useful to uncover relevant aspects for assessing cognitive functions as previously demonstrated for other modeling approaches. For example, parameters of drift-diffusion models of reaction times (Smith & Ratcliff, 2009) were found to be related to general intelligence (van Ravenzwaaij et al., 2011) and working memory (Schmiedek et al., 2007). Furthermore, distributional reaction time analysis may categorize healthy participants and patients suffering from psychiatric disorders (Kaiser et al., 2019; Karalunas et al., 2014; Vinogradov et al., 1998). The Rescorla-Wagner model could also be used for such differentiations, especially in the domain of belief-updating (Mengotti et al., 2017). By modeling a single cognitive process, however, the Rescorla-Wagner model is very dependent on the presence and size of a participant’s validity effects (see analysis in S3, showing that the correlation between model fit and cue-validity are higher for Rescorla-Wagner than bDCM).

BDCM, in contrast, simulated smoother reaction time distributions (larger number of non-significant p-values in KS-test), possibly providing a richer representation of the underlying processes. Although bDCM may reflect a portion of variance in the reaction time data that is not task-related, this variance could reflect the processes of belief-updating in a more complex brain-dynamics-dependent matter. BDCM is a *model of brain dynamics* that can, in principle, be applied to any task, while the Rescorla-Wagner model represents a specialized *model of a cognitive process*.

As bDCM can be applied to model different behavioral read-outs in various tasks, it can enhance our understanding of how DCM’s connectivity parameters relate to behavior. So far, this link could only be established using indirect methods, such as correlations between DCM parameters and behavioral measures across participants. For example, DCM’s task connectivity parameters have been related to symptoms of depression and schizophrenia (Desseilles et al., 2011; Schlösser et al., 2008; Wu et al., 2014), and have been correlated with behavioral measures before and after interventions using non-invasive neurostimulation (Grefkes et al., 2010). Although the investigation of such associations does not allow causal interpretations, bDCM enables more firm conclusions how brain dynamics in selected brain regions impact behavior.

Furthermore, brain and behavioral dynamics both regularize bDCM, so that the model parameters encode the most reliable set of information from both sources (Rigoux & Daunizeau, 2015). This procedure could yield more robust and stable connectivity estimates but also encode more specific information. This may be particularly relevant for so-called “generative embedding” approaches, where a generative model and its estimated parameters are used as a form of dimensionality reduction (Brodersen, Haiss, et al., 2011; Brodersen, Schofield, et al., 2011). In fact, this was confirmed by our findings, where it was possible to differentiate the horizontal from the vertical run only when using the bDCM model’s connectivity parameters. This makes bDCM a unique approach for the identification of biomarkers that are relevant for certain behaviors – provided that they are stable across participants and sessions (Elliott et al., 2020).

Since bDCM is a generative model, it can also be used to simulate how alterations to the underlying brain network might change behavior (Rigoux & Daunizeau, 2015). This allows simulating the behavioral effects of neuromodulatory interventions and the generation of new hypotheses and experiments. The guidance and information of computational models will eventually lead to a better understanding of the neural mechanisms underlying behavioral outcomes (Kriegeskorte & Douglas, 2018; Turner et al., 2017).

Unfortunately, applying artificial lesions to the network model in our study revealed technical problems of this approach. More specifically, the estimated models lacked numerical stability and required manual intervention, which substantially changed the model’s output. Even though some of the resulting patterns were consistent with the literature (e.g., an increase of the contralesional validity effect after a lesion to right TPJ and to a lesser extent in left TPJ (Malherbe et al., 2018; Posner et al., 1984)), other simulations were highly variable. Hence, the relatively novel bDCM approach’s potential problems, such as over-fitting and non-generalizability, need to be considered in future studies.

### Conclusion

bDCM was applied for the first time to reaction time data of a larger sample of participants. Our findings provided evidence for a considerable additional value of the method compared to a purely behavioral model and classical DCM and identified practical use issues. Data suggest that bDCM is indeed a promising tool to enhance our understanding of how brain dynamics generate specific behavioral patterns.

## Acknowledgements

SV was supported by funding from the Federal Ministry of Education and Research (BMBF, 01GQ1401). We also like to thank our colleagues at the INM-3 for their valuable feedback and suggestions throughout the development of the study and analyses.

## Data Availability Statement

The data that support the findings of this study are available on request from the corresponding author. The data are not publicly available due to privacy or ethical restrictions.

