## Supplemental Materials for "Simultaneous Modeling of Reaction Times and Brain Dynamics in a Spatial Cuing Task"

### S1 Details for fMRIprep

The preprocessing of functional and anatomical data was performed using FMRIPREP version 1.1.1 (Esteban et al., 2018, 2019, RRID:SCR_016216), a Nipype (RRID:SCR_002502, Gorgolewski et al., 2011, 2017) based tool, run as a docker-image. Each T1-weighted volume (T1w) was corrected for intensity non-uniformity using N4BiasFieldCorrection v2.1.0 (Tustison et al., 2010) and skull-stripped using antsBrainExtraction.sh v2.1.0 (using the OASIS template). Spatial normalization to the ICBM 152 Nonlinear Asymmetrical template version 2009c (Fonov, Evans, McKinstry, Almli, & Collins, 2009, RRID:SCR_008796) was performed through nonlinear registration with the antsRegistration tool of ANTs v2.1.0 (Avants, Epstein, Grossman, & Gee, 2008, RRID:SCR_004757), using brain-extracted versions of both T1w volume and template. Brain tissue segmentation of cerebrospinal fluid (CSF), white matter (WM) and gray matter (GM) was performed on the brain-extracted T1w using fast (Zhang, Brady, & Smith, 2001, FSL v5.0.9, RRID:SCR_002823).

Functional data were slice-time corrected using 3dTshift from AFNI v16.2.07 (Cox, 1996, RRID:SCR_005927) and motion-corrected using mcflirt (FSL v5.0.9, Jenkinson, Bannister, Brady, & Smith, 2002). "Fieldmap-less" distortion correction was performed by co-registering the functional image to the same-subject T1w image with intensity inverted (Wang et al., 2017) constrained with an average fieldmap template (Treiber et al., 2016), implemented with antsRegistration (ANTs). This was followed by co-registration to the corresponding T1w using boundary-based registration (Greve & Fischl, 2009) with 9 degrees of freedom, using flirt (FSL). Motion correcting transformations, field distortion correcting warp, BOLD-to-T1w transformation, and T1w-to-template (MNI) warp were concatenated and applied in a single step using antsApplyTransforms (ANTs v2.1.0) using Lanczos interpolation.

Frame-wise displacement (Power et al., 2014) was calculated for each functional run using the implementation of Nipype.

Many internal operations of FMRIPREP use Nilearn (Abraham et al., 2014, RRID:SCR_001362), principally within the BOLD-processing workflow. For more details of the pipeline see http://fmriprep.readthedocs.io/en/1.1.1/workflows.html.

### S2 fMRI – GLM analysis coordinates

Table 1 and Table 2 for the peak coordinates shown in Figure 4 of the main manuscript. The contrast invalid > valid is shown for each run of the fMRI experiment separately. To label the regions, we used the Harvard-Oxford atlas and reported the region with maximum probability.

Table S1: Peak statistics and clusters sizes for the horizontal T-map. Labels were automatically extracted from the Harvard-Oxford atlas, reporting the region with the highest probability from the coordinates.

**Horizontal Run**

| Cluster ID | X | Y | Z | Peak Stat (T) | Cluster Size (mm3) | Label |
|---|---|---|---|---|---|---|
| 1 | -27.25 | 8.62 | 57.3 | 5.57 | 8056 | Superior Frontal Gyrus |
| 1a | -46 | 5.5 | 37.5 | 4.43 | | Inferior Frontal Gyrus, pars opercularis |
| 1b | -24.12 | 2.38 | 73.8 | 3.85 | | Insular Cortex |
| 1c | -39.75 | -3.88 | 47.4 | 3.68 | | Inferior Frontal Gyrus, pars opercularis |
| 2 | 29 | 2.38 | 54 | 5.56 | 4350 | Superior Frontal Gyrus |
| 2a | 22.75 | 14.88 | 54 | 4.51 | | Insular Cortex |
| 3 | 7.12 | -69.5 | 57.3 | 5.4 | 15565 | Cingulate Gyrus, posterior division |
| 3a | -14.75 | -75.75 | 57.3 | 4.77 | | Angular Gyrus |
| 3b | -33.5 | -75.75 | 34.2 | 4.57 | | Angular Gyrus |
| 3c | -8.5 | -60.12 | 47.4 | 4.41 | | Cingulate Gyrus, posterior division |
| 4 | -49.12 | 21.12 | 27.6 | 5.11 | 1998 | Inferior Frontal Gyrus, pars triangularis |
| 5 | -33.5 | -53.88 | 37.5 | 5.08 | 6123 | Postcentral Gyrus, Supramarginal Gyrus, posterior division |
| 5a | -42.88 | -50.75 | 47.4 | 5.07 | | Supramarginal Gyrus, posterior division |
| 5b | -30.38 | -44.5 | 37.5 | 4.07 | | Supramarginal Gyrus, anterior division |
| 6 | -5.38 | 14.88 | 54 | 4.73 | 3222 | Subcallosal Cortex |
| 6a | 7.12 | 24.25 | 54 | 4.11 | | Insular Cortex |
| 7 | 41.5 | 27.38 | 24.3 | 4.5 | 3609 | Superior Frontal Gyrus |
| 7a | 44.62 | 8.62 | 34.2 | 4.26 | | Inferior Frontal Gyrus, pars opercularis |
| 7b | 50.88 | 24.25 | 40.8 | 3.93 | | Superior Frontal Gyrus |

Table S2: Peak statistics and clusters sizes for the vertical T-map. Labels were automatically extracted from the Harvard-Oxford atlas, reporting the region with the highest probability from the coordinates.

**Vertical Run**

| Cluster ID | X | Y | Z | Peak Stat (T) | Cluster Size (mm3) | Label |
|---|---|---|---|---|---|---|
| 1 | 60.25 | -53.88 | 17.7 | 7.18 | 7927 | Supramarginal Gyrus, posterior division |
| 1a | 54 | -41.38 | -2.1 | 5.62 | | Middle Temporal Gyrus, posterior division |
| 1b | 63.38 | -38.25 | -8.7 | 5.5 | | Middle Temporal Gyrus, posterior division |
| 2 | 41.5 | 27.38 | 14.4 | 6.29 | 21269 | Middle Frontal Gyrus |
| 2a | 44.62 | 5.5 | 30.9 | 5.81 | | Inferior Frontal Gyrus, pars opercularis |
| 2b | 41.5 | 33.62 | 24.3 | 5.01 | | Superior Frontal Gyrus |
| 2c | 29 | -0.75 | 54 | 5 | | Superior Frontal Gyrus |
| 3 | 7.12 | 21.12 | 54 | 6.09 | 4898 | Insular Cortex |

| | | | | | | |
|---|---|---|---|---|---|---|
| 3a | -5.38 | 11.75 | 57.3 | 4.02 | | Insular Cortex |
| 4 | -36.62 | -0.75 | 40.8 | 6 | 11053 | Inferior Frontal Gyrus, pars opercularis |
| 4a | -24.12 | -13.25 | 54 | 5.02 | | Inferior Frontal Gyrus, pars opercularis |
| 4b | -33.5 | -3.88 | 57.3 | 4.84 | | Superior Frontal Gyrus |
| 4c | -49.12 | 24.25 | 30.9 | 4.75 | | Superior Frontal Gyrus |
| 5 | 38.38 | -57 | 57.3 | 5.39 | 33612 | Angular Gyrus |
| 5a | -36.62 | -53.88 | 47.4 | 5.23 | | Postcentral Gyrus |
| 5b | 19.62 | -72.62 | 57.3 | 5.19 | | Angular Gyrus |
| 5c | 35.25 | -60.12 | 44.1 | 5.19 | | Angular Gyrus |
| 6 | -39.75 | 21.12 | -5.4 | 5.18 | 4382 | Frontal Pole |
| 6a | -33.5 | 24.25 | 1.2 | 4.89 | | Frontal Pole |
| 6b | -27.25 | 18 | -15.3 | 4.57 | | Cuneal Cortex |
| 6c | -42.88 | 24.25 | 11.1 | 4.29 | | Middle Frontal Gyrus |

### S3 Correlation between fit-statistic and validity differences

To test whether the size of the validity effect influenced the resulting fit-statistic, we correlated (using Pearson's correlation) each permutation of error ($R^2$, mean absolute error), model (Rescorla-Wagner), and run with the validity difference (mean RT invalid – mean RT valid). The $R^2$-score was highly correlated with the validity difference, where the effect appeared to be stronger for the Rescorla-Wagner model than for the behavioral DCM. There was, however, no correlation with the mean absolute error (MAE).

Table S3: Pearson's Correlations between the validity difference and the different fit-statistics.

| **Model** | **Score** | **Run** | **r** | **adj_r2** | **p-value** | **$BF_{10}$** |
|---|---|---|---|---|---|---|
| Rescorla Wagner | MAE | Horizontal | 0.214 | -0.037 | 0.294 | 0.410 |
| | | Vertical | 0.112 | -0.073 | 0.586 | 0.28 |
| | $R^2$ | Horizontal | 0.817 | 0.639 | < 0.001 | $5.06 * 10^7$ |
| | | Vertical | 0.854 | 0.706 | < 0.001 | $4.86 * 10^8$ |
| BDCM | MAE | Horizontal | 0.232 | -0.028 | 0.253 | 0.452 |
| | | Vertical | 0.108 | -0.074 | 0.600 | 0.277 |
| | $R^2$ | Horizontal | 0.719 | 0.475 | < 0.001 | 795.149 |
| | | Vertical | 0.789 | 0.589 | < 0.001 | $1.22 * 10^7$ |

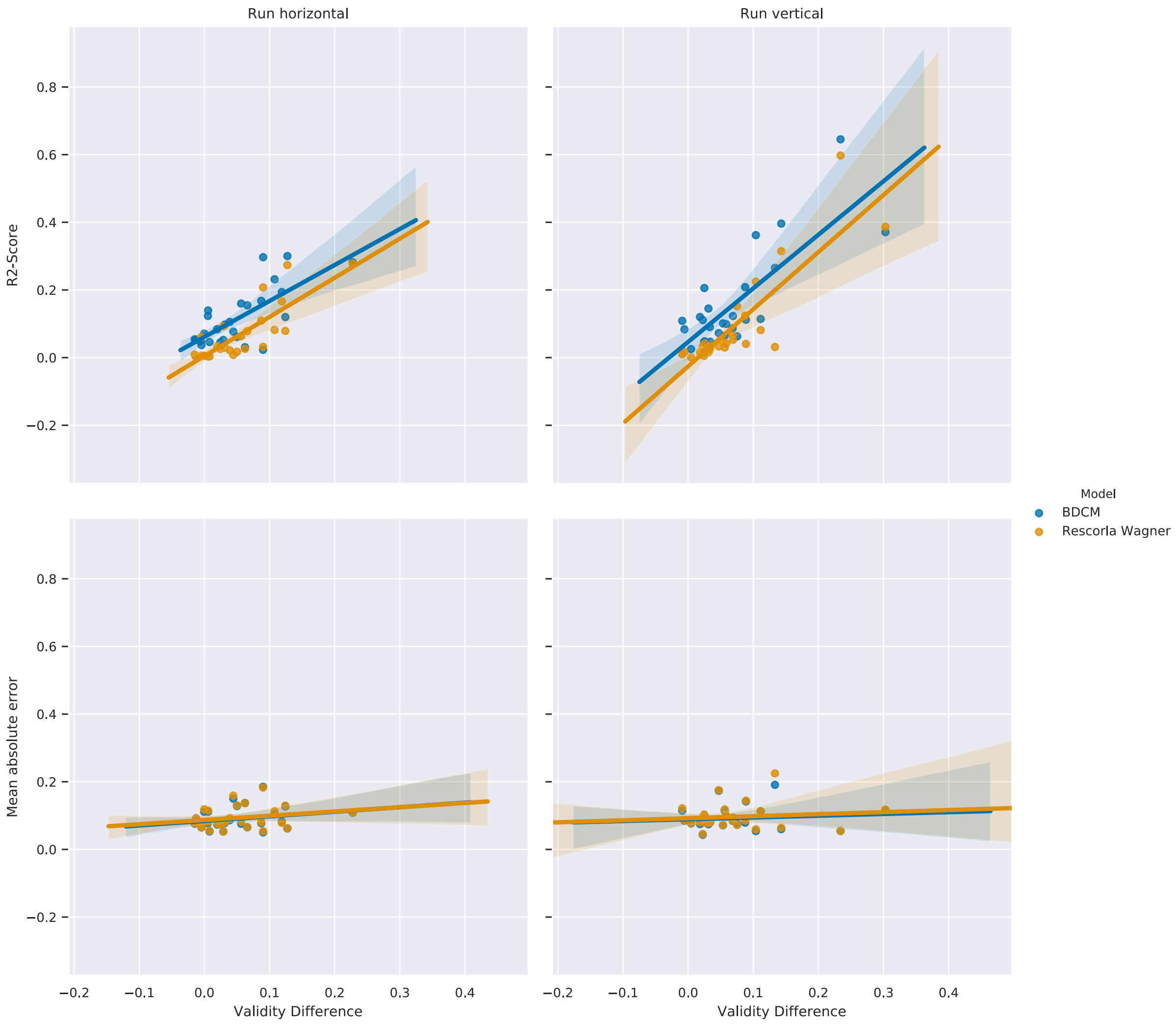


Figure S1: Pearson correlation between the validity difference (y-axis) and the fit-statistics (x-axis) for the two models, the different metrics (rows), and the two runs (columns).

### S4 Lesion analysis additional reporting

We used a similar approach as before to compare the fit-statistics between the data with an increased self-inhibition and the regular models. The dampened model performed significantly worse than the original models, with the R2-score being particularly affected, while the mean absolute error remained relatively stable. This analysis showed why we ought to be cautious in the analysis of the lesion data results.

Figure S2 is the analog to Figure 9 in the main manuscript, for lesions simulated for up-wards and down-wards orienting.

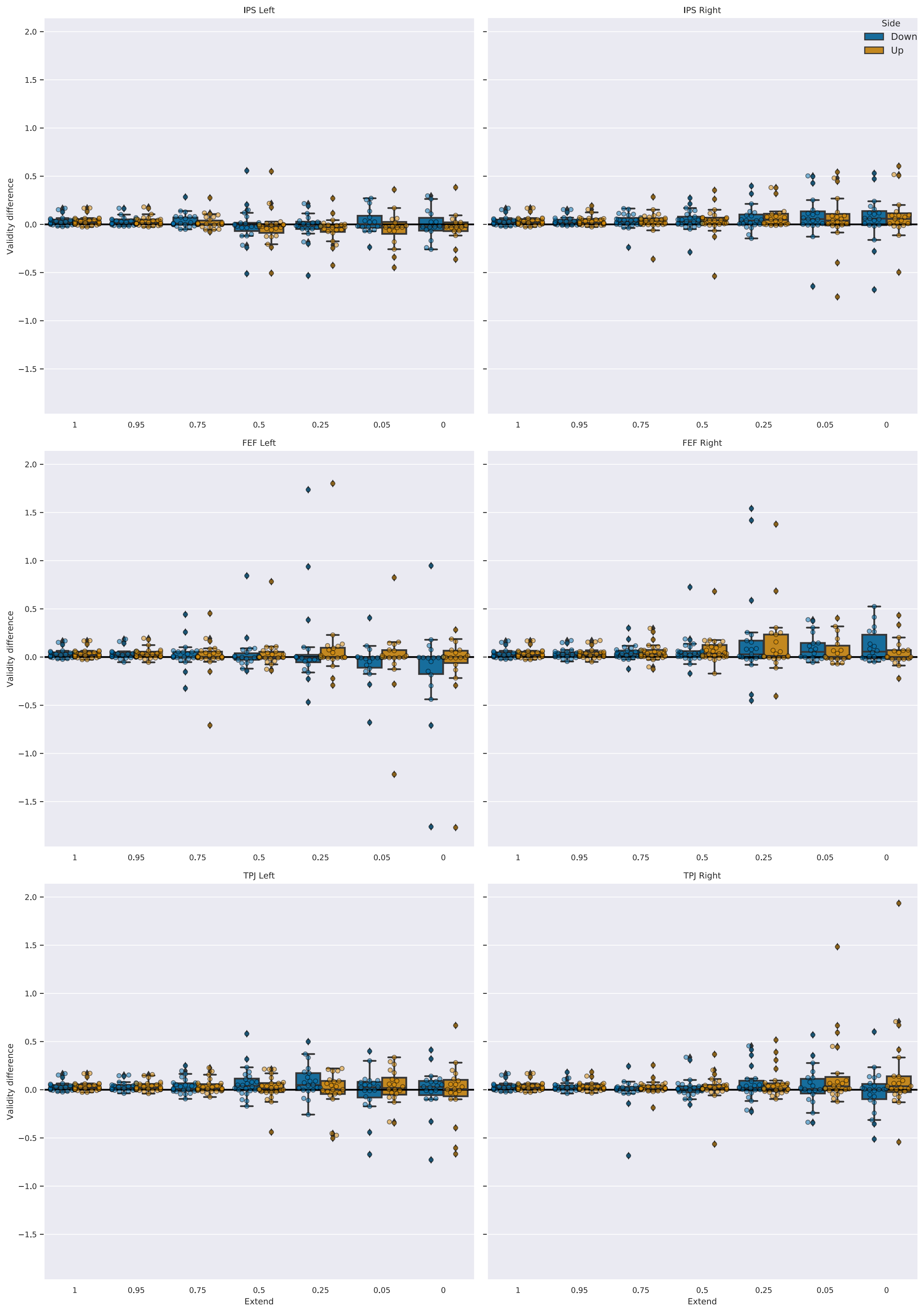


Figure S2: Validity effect for simulated reaction times for the vertical run, after brain lesions of different extent. An extent of 1 means that no lesion occurred, an extent of 0 indicates that the region was fully disabled. Boxplots indicate the median of the data, the IQR, and the minimum and maximum values. Outliers exceeded the 1.5 * IQR criterion.

### S5 Connectivity matrices

Figures S3-S6 displays the average connectivity strength of the (behavioral) dynamic causal modeling analysis. Connectivity parameters were averaged across participants. Inhibitory connections (A matrix diagonal) were exponentiated before averaging.

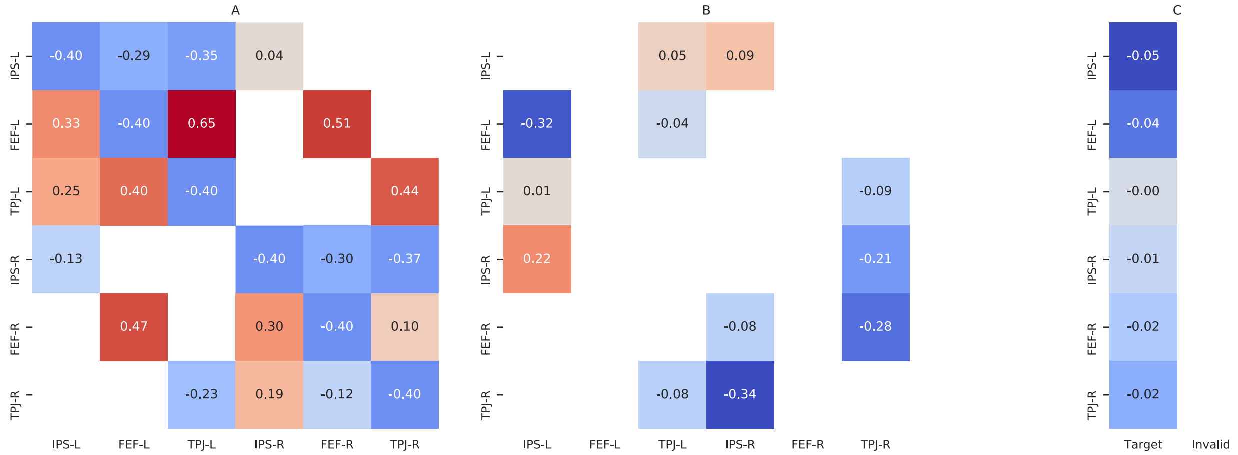


Figure S3: The average strength of the DCM connectivity parameters of the horizontal run.

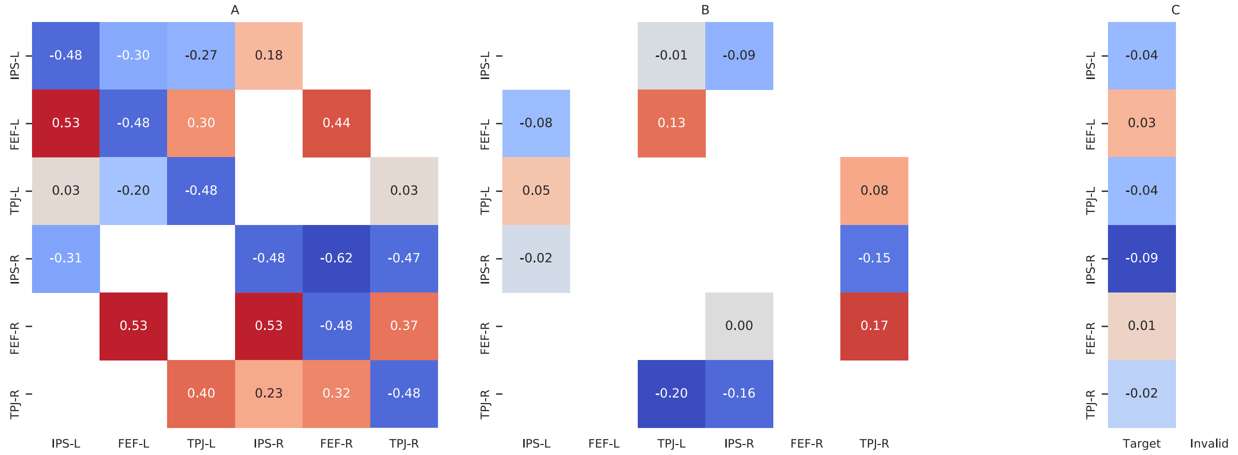


Figure S4: The average strength of the DCM connectivity parameters of the vertical run.

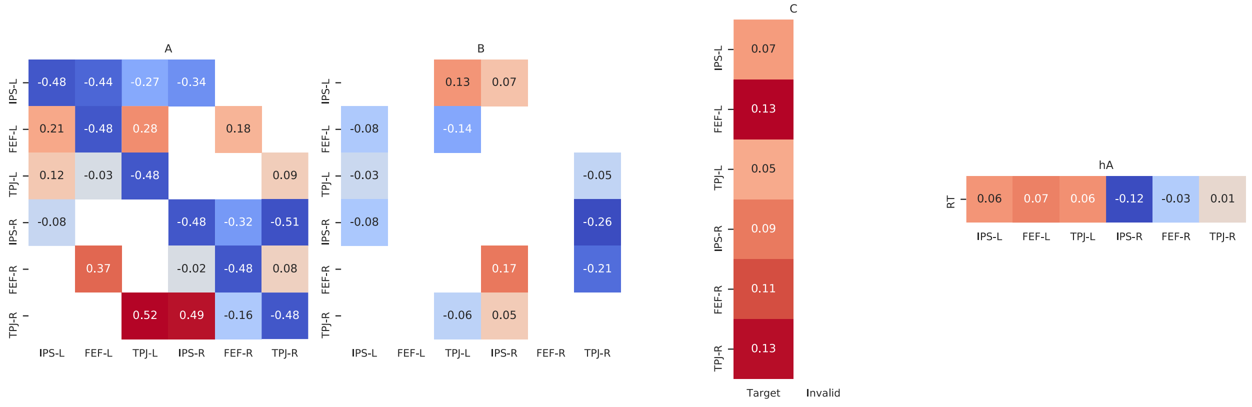


Figure S5: The average strength of the bDCM connectivity parameters of the horizontal run.

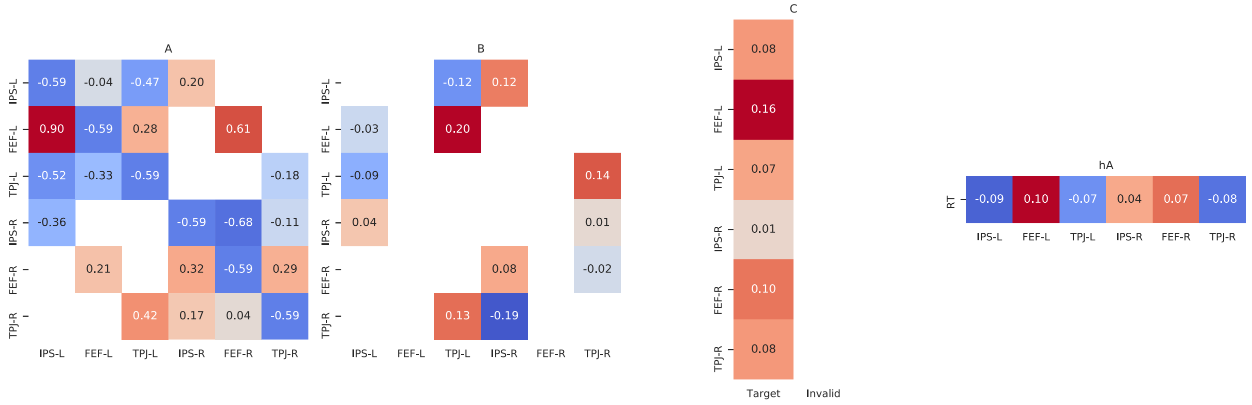


Figure S6: The average strength of the bDCM connectivity parameters of the vertical run.
